# A coregulatory influence map of glioblastoma heterogeneity and plasticity – From cells in vitro to tumors and back again

**DOI:** 10.1101/2024.08.23.609303

**Authors:** Chloé Bernhard, Konstantinos Geles, Geoffrey Pawlak, Wajdi Dhifli, Aurélien Dispot, Jules Dusol, Maria Kondratova, Sophie Martin, Mélissa Messé, Damien Reita, David Tulasne, Isabelle Van Seuningen, Natacha Entz-Werle, Silvia Anna Ciafrè, Monique Dontenwill, Mohamed Elati

**Affiliations:** UMR7021 CNRS, University of Strasbourg, Illkirch, France; Univ. Lille, CNRS, Inserm, CHU Lille, UMR9020-U1277 – CANTHER – Cancer Heterogeneity Plasticity and Resistance to Therapies, Lille F-59000, France; Dept. of Cancer Molecular Genetics, Laboratory of Biochemistry and Molecular Biology, University Hospital of Strasbourg, 67200 Strasbourg, France; Pediatric Onco-Hematology Unit, University Hospital of Strasbourg, 67098 Strasbourg, France; Dept. of Biomedicine and Prevention, University of Rome “Tor Vergata”, Rome, Italy

**Keywords:** GBM-cRegMap, glioblastoma, coregulatory network, heterogeneity, plasticity, reference mapping, machine learning

## Abstract

We present GBM-cRegMap, an online resource providing a comprehensive coregulatory influence network perspective on glioblastoma (GBM) heterogeneity and plasticity. Using representation learning algorithms, we derived two components of this resource: GBM-CoRegNet, a highly specific coregulatory network of tumor cells, and GBM-CoRegMap, a unified network influence map based on 1612 tumors from 16 studies. As a widely applicable closed-loop system connecting cellular models and tumors, GBM-cRegMap will provide the GBM research community with an easy-to-use web tool (https://gbm.cregmap.com) that maps any existing or newly generated transcriptomic “query” data to a reference coregulatory network and a large-scale manifold of disease heterogeneity. Using GBM-cRegMap, we demonstrated the synergy between the two components by refining the molecular classification of GBM, identifying potential key regulators, and aligning the transcriptional profiles of tumors and in vitro models. Through the amalgamation of a vast dataset, we validated the proneural (PN)-mesenchymal (MES) axis and identified three subclasses of classical (CL) tumors: astrocyte-like (CL-A), epithelial basal-like (CL-B), and cilium-rich (CL-C). We revealed the CL-C subclass, an intermediate state demonstrating the plasticity of GBM cells along the PN-MES axis under chemotherapy. We identified key regulators, such as PAX8, and NKX2.5, involved in TMZ resistance. Notably, NKX2.5, more expressed in higher-grade gliomas, negatively impacts patient survival and regulates genes involved in glucose metabolism.

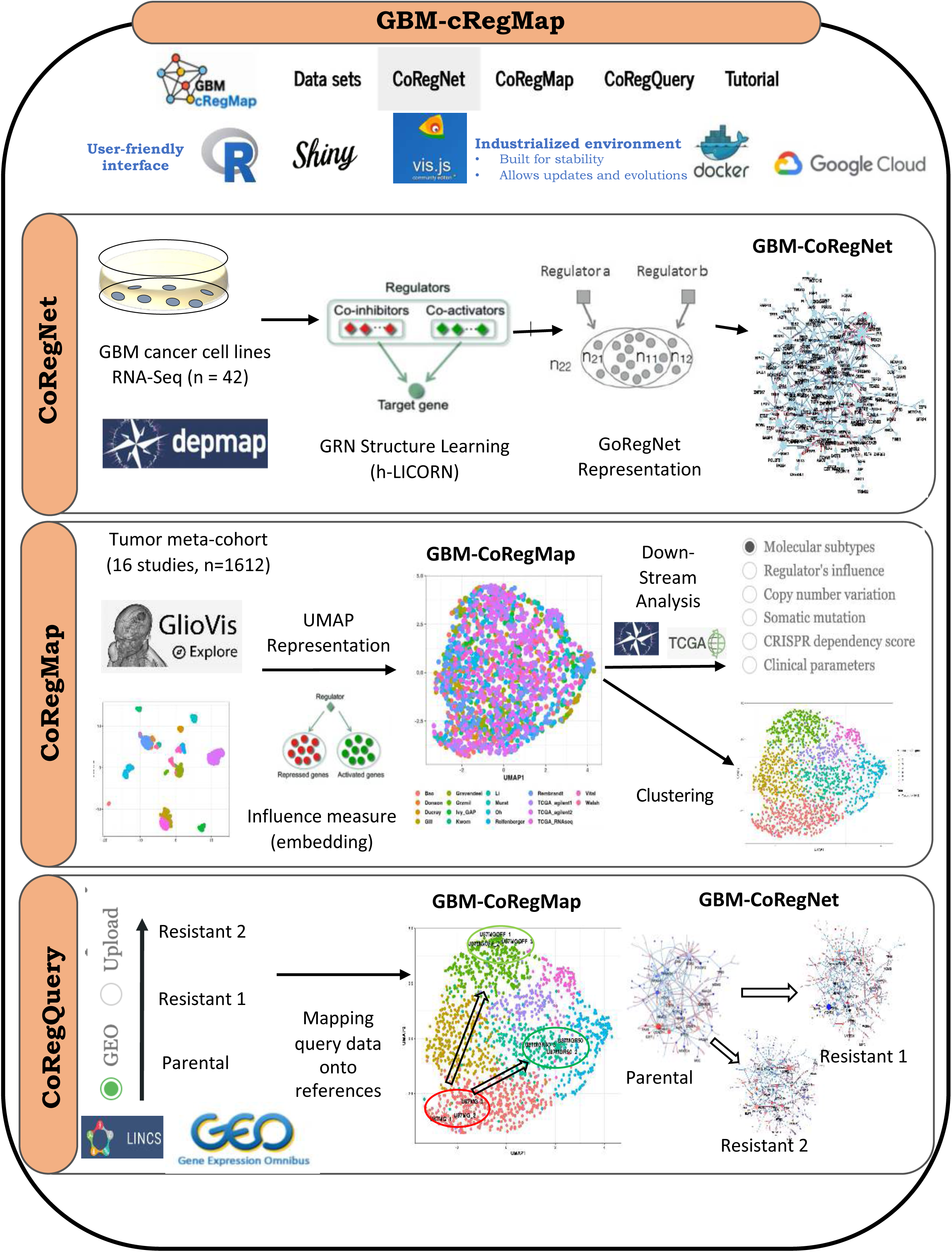

## Introduction

Glioblastoma (GBM), the most common and aggressive tumor of the brain and central nervous system, represents a major challenge in neuro-oncology. According to the CBTRUS data repository^1^, GBM accounts for 48.6% of malignant brain tumors and 57.7% of all gliomas, and is one of the most lethal and aggressive forms of cancer. Standard treatment involves surgery, chemotherapy and radiotherapy (Stupp protocol), but even with this treatment, most patients succumb within 15–18 months^1^. GBMs are intrinsically highly heterogeneous in terms of the histological and molecular characteristics.

Previous studies attempting to classify GBM into molecular subtypes based on bulk gene expression profiling have met with various degrees of success. The Philips group first defined three major tumor subsets: proneural, mesenchymal, and proliferative^2^. Verhaak *et al*.^3^ sub-sequently proposed a classification into four subtypes: proneural (PN), neural (NEU), classical (CL), and mesenchymal (MES). This classification was later revised, with exclusion of the neural subtype^4^. More recent single-cell studies have revealed additional layers of heterogeneity due to the plasticity of transcriptional programs within the same tumor and the dominating metabolic pathway. Neftel *et al*.^5^ deconvoluted the phenotypic states of GBM cells into four major line-age-specific cellular identities: astrocyte-like (AC), mesenchymal-like (MES), neural progeni-tor cell-like (NPC) and oligodendrocyte progenitor cell-like (OPC). The Iavarone group^6,7^ identified four states, based on the most active pathways, along two axes: a metabolic axis including a mitochondrial (MTC) and a glycolytic/plurimetabolic (GPM) state, and a neurodevelopmental axis including a proliferative/progenitor (PPR) and a neuronal (NEU) state.

These observations suggest that one of the most promising ways to dissect GBM heterogeneity is to analyze the gene regulatory networks (GRN) that give rise to different molecular subtypes and cell states. The computational approaches that have proved successful in this field to date are those that allow the reverse-engineered construction of context-specific GRNs (e.g., ARACNe^8^ and LICORN^9,10^ using a bulk-transcriptome, and SCENIC^11^ using single-cell data). A second key breakthrough in regulatory network exploration made possible by CoReg-Net^12,13^ and VIPER^14^ was the consideration of regulator activity, rather than just the production of regulators, based on evaluations of the expression of target genes with the aim of detecting master regulators. Several recent studies on GBM have inferred PN/MES subtype-specific regulatory networks, mostly from TCGA patient or single-cell data. Three important transcription factors have been identified by SCENIC: *FOSL2, CEBPB*, and *EPAS1*, which predominate in mesenchymal cells^15^. Similarly, previous studies with ARACNE confirmed that CEBPB and STAT3, the master regulators, cooperate with *FOSL2* to mediate the PN to MES transition^16^.

The application of -omics and systems biology approaches has, therefore, made a significant contribution to GBM research. However, the field now faces the challenge of dealing with data and knowledge distributed among a large number of studies and databases. Each study has entailed a marked increase in data size, complexity, and specificity. Consequently, fostering collaboration among biologists requires concerted efforts to centralize the data from these investigations into an intuitive system-level shared model. We have addressed this challenge by developing GBM-cRegMap, a robust web-based tool providing researchers with rapid access to a unified coregulatory influence network view of GBM. GBM-cRegMap delves into representation learning, a pivotal aspect of machine learning in which algorithms extract significant patterns from raw data and transform them into more interpretable, accessible, and shared representations. The GBM-cRegMap tool incorporates the workflow illustrated in the graphical abstract, integrating data from over 20 transcriptomic studies of tumors and cell lines (Supplementary Table S1). The GBM-cRegMap tool enables users to explore similarities and differences between subtypes, identify possible core regulators, detect rare subtypes, associate tumors with cell-line transcriptomic profiles, and define new targets associated with different tumor states and plasticity. GBM-cRegMap boasts an intuitive user interface (UI) that is easy to use, even for those with no expertise in bioinformatics. For all the networks and plots generated, users can run various annotations, add new data, and download the raw data required to generate plots for future analyses or publications. The wide range of computational network biology, machine learning, visualization, bioinformatics software and other resources required to achieve this are listed in Supplementary Table S2.

We demonstrated the utility of GBM-cRegMap by proposing an alternative description of GBM cancer heterogeneity based on regulatory network activities detectable across cohorts. Through the amalgamation of a vast dataset, we validated the proneural (PN)-mesenchymal (MES) axis and identified three subclasses of classical (CL) tumors: astrocyte-like (CL-A), epithelial basal-like (CL-B), and cilium-rich (CL-C). We also identified a neural normal-like subclass (NL). In addition, we identified new targets associated with different tumor states, plasticity, and resistance to therapy, showcasing the versatility and significance of GBM-cRegMap for advancing our understanding of GBM at the system level.

## Results

### Representation Learning and Evaluation of GBM-cRegMap Reference Components

We designed the GBM-cRegMap resource around two synergistic reference compounds: GBM-CoRegNet, a highly specific coregulatory network of tumor cells, and GBM-CoRegMap, a unified network influence map based on 1,612 tumors from 16 studies. To generate these reference compounds, we first used h-LICORN^9^ to infer a gene regulatory network (GRN) from the transcriptomes of 42 GBM cancer cell lines from CCLE 2019Q1 with a curated list of regulators composed of transcription factors (TFs) and co-factors (co-TFs) (*n* = 2,375). We used transcriptomes from GBM tumor-derived cell lines to strengthen the signal specific to GBM cells and to prevent the introduction of possible bias due to the use of less homogeneous transcriptomic data from GBM tumors which would include non-specific signals and signals associated with the tumor microenvironment. The resulting GRN was composed of 539 coregulators (Supplementary Table S3), 8269 target genes and 32360 regulatory interactions, with a significant enrichment in validated TF-gene interactions and transcription factor-binding sites (*p* <1e-100). Based on the shared targets of each pair of TFs/co-TFs (Supplementary Table S4), the GBM-GRN was transformed into a coregulatory network (GBM-CoRegNet) with the CoRegNet package^13^. GBM-CoRegNet (Fig. 1A) contains 2171 cooperative interactions enriched in protein-protein interactions (PPIs; *p* <1e-100).

**Figure 1.**
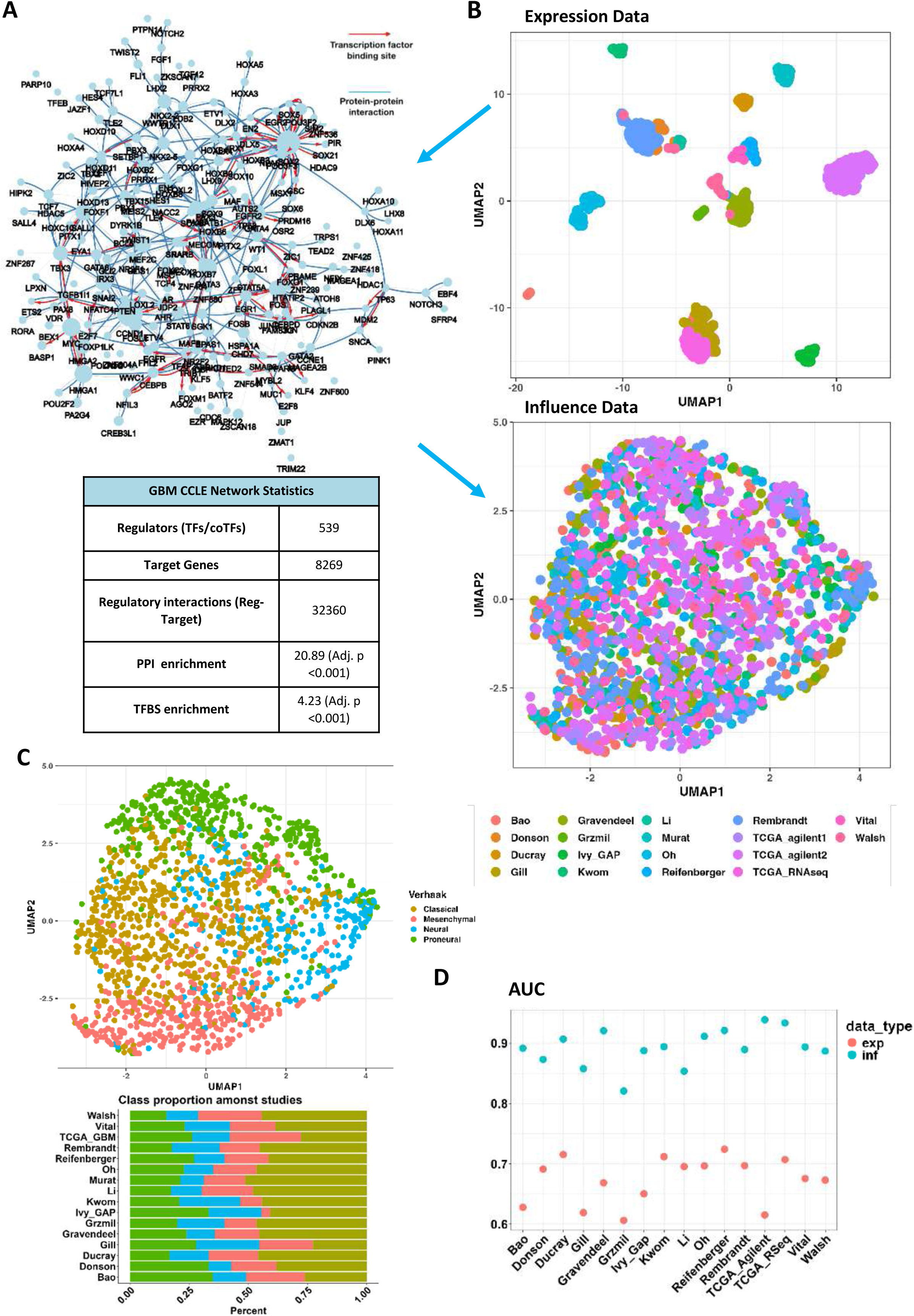
Evaluation of GBM-cRegMap reference components. (A) Graphical representation (top) of the co-operativity network inferred from the transcriptome of 42 GBM cancer-derived cell lines (GBM-CoRegNet). Nodes represent transcription factors and cofactors (TFs/co-TFs). Node size is proportional to the number of targets of the TF/co-TF. Coregulatory interactions between nodes are indicated: protein-protein interactions with published evidence are shown in blue, transcriptional regulation interactions with published evidence are shown in red, and interactions defined solely by the h-LICORN algorithm are shown in gray. The table (bottom) summarize metrics from the GBM-CoRegNet. (B) UMAP visualization of the metacohort of 1629 tumors, illustrating the difference between the use of transcriptomic (top) and influence (bottom) data from different datasets. Influence data produce a batch effect-free metacohort from the original RNA expression values. (C) UMAP visualization of the GBM-cRegMap metacohort, using sample annotation colors derived from the Verhaak classification. (D) The Area Under the Curve (AUC) for the resulted classifier of Verhaak subtypes using gene expression or influence in cross-batch prediction. This cross-batch prediction involves training the classifier on one dataset and testing it on others. Used classifier: PAMR (prediction analysis for microarrays).

As for the source of GBM-CoRegMap, we compiled a GBM metacohort from 16 GBM tumor cohorts (Supplementary Table S1) and calculated sample-independent gene regulatory influences with the inferred GBM-GRN (see Methods)^13^. We used minimalistic preprocessing pipelines for microarrays or RNA-sequencing data to achieve transparency and preprocessing parity (See Methods). The resulting metacohort influence data yielded both significant feature reduction and sample size augmentation (single GBM bulk expression cohort: <500 samples x ≃18000 features vs. meta-cohort influence data: 1612 samples × 539 features). Based on its greater sensitivity, and its ability to capture both local and global relationships^17^, we chose to use Uniform Manifold Approximation and Projection (UMAP)^18^ to visualize the meta-cohort influence data in a two-dimensional embedded space, hereafter denoted “GBM-CoRegMap”. We ensured that the resulting GBM-CoRegMap simulated tumor heterogeneity and did not result from potential confounding factors, by annotating the samples with Verhaak subtypes according to the corresponding published gene expression classifier. Using influence-based embedding, GBM-CoRegMap successfully reproduced a manifold that are not driven by batch effects, compared to the original expression space (Fig. 1B) and aligns with the known heterogeneity of GBM (Fig. 1C). Furthermore, relative to gene expression, this influence-based Verhaak subtype classifier yielded a mean 33% increase in cross-batch prediction performance, as evaluated by calculating the area under the curve (AUC) with the PAMR (prediction analysis for microarrays) classifier (Fig. 2D).

**Figure 2.**
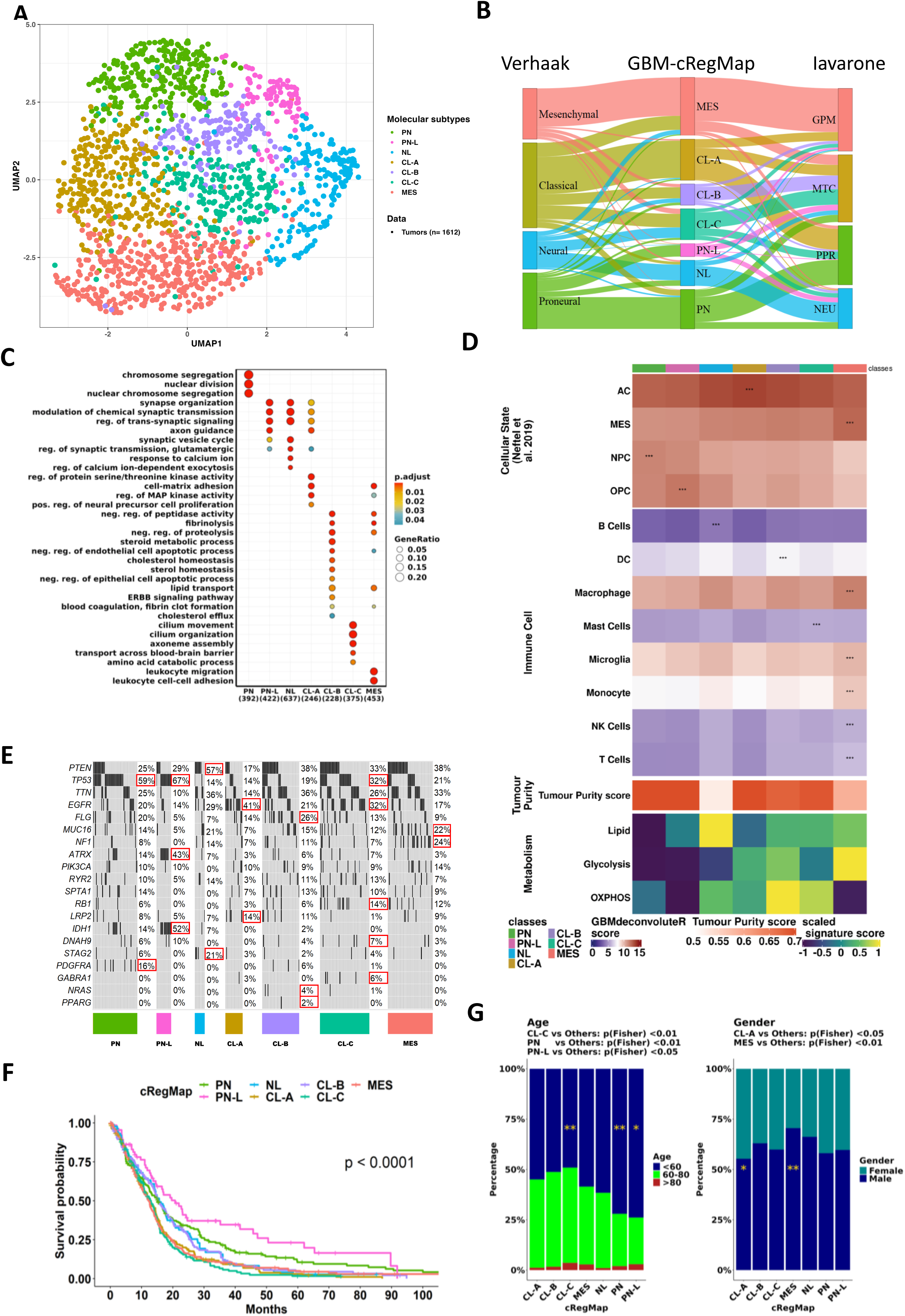
Molecular and clinical characteristics of GBM-cRegMap classes. (A) UMAP visualization of the GBM-cRegMap metacohort, using sample annotation colors derived from the GBM-cRegMap classification. (B) Alluvial plot displaying the GBM-cRegMap metacohort samples with state-of-the-art classifications of glioblastoma (Verhaak^3^ and Iavarone^7^ classifications on the left and right, respectively) and the corresponding classification proposed by GBM-cRegMap (in the middle). (C) Dot plot with significantly enriched Gene Ontology biological processes for each GBM-cRegMap class derived from the clusterProfiler R package. GO/BPs with an adjusted *p*-value <0.05 were considered; *p*-values were adjusted by the Benjamini-Hochberg method. (D) Heatmap displaying the deconvolution of cellular component scores with the GBMdeconvoluteR tool and metabolic gene signature scores with the sing-score R package. Asterisks indicate the highest score per row. (E) Oncoplot showing the most important genomic alterations associated with cRegMap classes, based on available data from the TCGA-GBM dataset. Red squares indicate the highest rates of alteration per gene for each class. (F) Overall survival curve stratified by cRegMap class. Survival analysis was performed by the Kaplan-Meier method, with the R package survival. Kaplan-Meier curves were generated from the available metacohort clinical data for 1110 patients. (G) Gender and age differences in GBM-cRegMap subclasses.

### Reference-mapping and UI Features of GBM-cRegMap

Gene expression profiling is routinely used by the GBM community to understand the disease. However, interpreting processed data to gain insights into biological mechanisms remains challenging, particularly for non-bioinformaticians. Many data visualization and network analysis tools require extensive data formatting, parameter tuning, significant running time, and are impractical with a limited number of samples. The concept of mapping newly generated biological “query” data to reliable references is a powerful idea that predates transcriptomic and network analysis. To advance this approach, we have created GBM-cRegMap (https://gbm.cregmap.com, http://github.com/i3bionet/gbm-cregmap), an open-source software package and web tool with GBM-CoRegNet and GBM-CoRegMap references and an interface for user-friendly analysis. The UI is composed of three functional components (see Graphical abstract): “CoRegNet”, “CoRegMap”, and “CoRegQuery”. The GBM-CoRegNet network is visualized via a Shiny applet in the R package visNetwork^19^. The CoRegNet network representations are intuitively understandable to biologists (Fig. 1A). They also facilitate the integration of several layers of information over nodes and edges. The CoRegNet tab allows users to investigate and evaluate the extent to which phenotypes of interest (e.g., tumor subtypes, wild-type vs. treated cell lines), each characterized by a set of active coregulators, colocalize within the reference GBM-CoRegNet network through an intuitive visualization. The second feature is GBM-CoRegMap, a unified influence map for 1612 tumors from 16 studies, enhanced by the inclusion of downstream analysis data, such as annotation of the map with molecular subtype, tumor purity, regulator influence, copy number variation, somatic mutations, clinical data, or gene essentiality scores. It can also be used for the alignment and colocalization of patient-derived cell models within the reference tumor heterogeneity map. Finally, query datasets can be mapped to GBM-cRegMap references with the “CoRegQuery” tab. GBM-cRegMap allows users to upload their expression datasets or to retrieve expression datasets from the Gene Expression Omnibus (GEO) with ease. As input, expression data can be uploaded in CSV format or as a GEO series ID, also known as a GEO accession code, from which GBM-cRegMap extracts the gene expression dataset and experimental information. GBM-cRegMap rapidly calculates regulator influences for new data, presenting the results in the GBM-CoRegNet compound and making it possible to investigate selected coregulators, interactions and systems further for relevance to the study. Furthermore, a supervised machine learning procedure (see Methods) increases the efficiency of the tool for predicting the localization of the query dataset samples within GBM-CoRegMap. This ensures consistent comparisons of tumor transcriptional patterns, facilitating potential explorations of tumor heterogeneity and the adaptability of the studied phenotypes.

### A Meta-Cohort Analysis of 1612 Tumors Refining the GBM Molecular Classification into Seven Classes of Biological and Clinical Relevance

Consistent with other publications on GBM^4,6^, the resulting metacohort influence data (1612 tumors × 539 regulators) show (Fig. 1C) that the MES/PN subtypes was the most homogeneous, whereas the CL and NEU subtypes were the least homogeneous (as assessed by lowest silhouette scores of 0.2 and 0.4 respectively). This highlights the opportunity for a refinement of the Verhaak classification. The large sample size of GBM-CoRegMap provides greater statistical power for subtype identification and minimizes sampling bias^20^.

GBM-cRegMap-based Seurat clustering (see Methods) resulted in seven distinct molecular classes (Fig. 2A). These classes were named on the basis of prior Verhaak subtyping (Fig. 2B). Sample size differed between classes, as follows: 17% proneural (PN), 6% proneural-low proliferative (PN-L), 11% neural normal-like (NL), 18% classical astrocyte-like (CL-A), 9% classical epithelial basal-like (CL-B), 14% classical cilium-rich (CL-C), and 25% mesenchymal (MES). The proportions of the seven GBM-cRegMap subtypes were almost identical in the different cohorts indicating the stability of the classification. We first characterized the GBM-cRegMap classes at both the transcriptomic (Fig. 2C-D, Supplementary Tables S6-7) and genomic (Fig. 2E) levels, to identify associated biological processes. We then assessed their clinical data and outcome (Fig. 2F-G).

The PN, PN-L, and NL subclasses were characterized principally by attributes of developmental processes, displaying proneural, gliogenesis, and neural signatures. The PN and PN-L subclasses accounted for the majority of PN-Verhaak tumors, indicating a further subdivision of the proneural subtype into two distinct entities. These two subclasses were also associated with progenitor cell states (NPC- and OPC-like, respectively), as described by Neftel *et al*.^5^. The PN subclass was enriched in cell division and mitotic cell cycle pathways and displayed the highest levels of expression for the proliferation markers *MKI67* and *DLGAP5*. By contrast, PN-L tumors had low levels of proliferation. Both subclasses had a higher frequency of *TP53* mutations (> 59%) than the other subtypes (0–32%). *PDGFRA* mutations were almost exclusive to the PN subclass (PN, 16%; PN-L, 0%), whereas *IDH1* mutations were more frequent in the PN-L subclass (PN-L, 52%; PN, 14%; NL, 7%; others, ∼0%). Notably, the 2021 WHO-CNS^21^ classification no longer classifies IDH-mutant gliomas as GBM. The NL subclass accounted for 42% and 49% of NEU_Verhaak and NEU_Iavarone subtype tumors, respectively. This subclass included a distinct cluster of GBM with a ‘normal-like’ transcriptomic profile enriched in neuron development-associated pathways and neuroglioma synaptic communication. We identified a common gene signature with the newly identified ‘normal-like’ IDH-WT subtype reported by Nguyen *et al*.^22^, which included genes such as *SLC32A1* (vesicular gamma-aminobutyric acid transporter) and *SYT5* (synaptotagmin 5). NL tumors harbored *PTEN* mutations in 57% of the cases, and show a higher frequency (21%) of *STAG2* mutations compared to other subclasses. This latter finding is intriguing, in view of the role of STAG2 as a tumor suppressor linked to therapeutic response to PARP inhibitors^23^. Moreover, STAG2 has been recently shown to modulate sensitivity to pharmacologic EZH2 inhibition in glioblastoma^24^.

The CL-A, CL-B, and CL-C subclasses accounted for the majority of CL_Verhaak tumors. Unlike the CL-A and CL-B subclasses, the CL-C subclass was evenly distributed across the CL and NEU Verhaak subtypes (Fig. 2B). Importantly, these three subclasses define previously unknown GBM subtypes based on tumor microenvironment and metabolic, mutational, and clinical information. The CL-A subclass was enriched in an astrocyte-like (AC) signature (Fig. 2D, Cellular State score^25^). It also displayed an upregulation of *MEOX2* homeobox TF and fatty-acid synthesis genes (*ELOVL2, ACSL3, PLA2G5*), and high levels of expression of the stem cell markers *NES* and *SOX9* (Supplementary Table S6). CL-B displayed enrichment in factors associated with stem cell survival (*SERPINB3*, *NANOG*, *SALL4*), the coagulation pathway, keratinization, and cell-cell adhesion (Fig. 2C) — features commonly expressed in basal squamous epithelial cells^26–28^. Notably, pseudo-epithelial and epithelial morphologies, although uncommon in GBM, are recognized as subtypes in the 2021 WHO classification of CNS tumors and are potentially associated with a poor prognosis^7^.

Furthermore, the enrichment of CL-B subclass in the oxidative phosphorylation (OXPHOS) pathway signature (Fig. 2D) and cholesterol homeostasis and metabolism of steroids (Fig. 2C, Supplementary Table S7) suggests an association with a mitochondrial subtype. This link is supported by shared metabolic intermediates and regulatory crosstalk between OXPHOS and cholesterol pathways in biological systems^29^. Notably, recent research underscores the critical role of cholesterol homeostasis in glioma stem cells^30^. Hence, targeting mitochondrial function and lysosomal cholesterol homeostasis could offer a promising therapeutic strategy for CL-B tumors, potentially leveraging specific mitochondrial inhibitors^31^.

Unlike CL-A and CL-C tumors, CL-B tumors did not over-express EGFR and had lower *EGFR* mutation rates (CL-A, 41%; CL-B, 21%; CL-C, 32%). By contrast, they displayed high rates of *FLG* mutation (CL-A, 14%; CL-B, 26%; CL-C, 13%) and were the only tumors in which infrequent *NRAS* (4%) and *PPARG* (2%) mutations are associated. Surprisingly, the CL-C (cilium-rich) subclass was found to be enriched in processes linked to cilia, including cilium assembly, movement, and organization, along with OXPHOS, lipid metabolism, and dopamine metabolism (Fig. 2C-D, Supplementary Table S7). The transcriptional landscape of CL-C tumors was further shaped by the activation of pivotal cell signaling pathways, mediated by *STAT3* and *RPS6KA1*, for example, and epigenetic regulators, such as *HDAC1, HDAC5, CBX4,* and *SUV39H2*. The CL-C subclass was also characterized by the upregulation of several prominent genes, including *DDIT4L*, *LGR6* (known to facilitate DNA repair and chemoresistance^32,33^), and *ETNPPL* (a negative regulator of glioma growth^34^). Unlike the other two classical subclasses, CL-C tumors had a weaker proliferative signature (*p* < 0.01) and higher rates of *TP53* (CL-A, 14%; CL-B, 19%; CL-C, 32%) and *DNAH9* (CL-C, 7%) mutation (Fig. 2E). Recent studies have linked suppressed ciliogenesis in glioma stem cells (GSCs) to continuous self-renewal, whereas the restoration of ciliogenesis shifts GSCs towards the differentiation state^35^. In addition, it has very recently been shown that primary cilia, through the regulation of the intracellular release of IL-6, contribute to GBM-driven immunosuppression^36^. Finally, the MES subclass was closely associated with the MES_Verhaak subtype and the single-cell MES-like state. This association reflects the features linked to a mesenchymal identity reported, such as higher mutation rates and low levels of *NF1* expression^34^. The MES subclass is enriched in the epithelial-mesenchymal transition (EMT) and immune-associated pathways. It also displayed an enrichment in glycolysis/hypoxia-related functions and lipid metabolism but no enrichment in the OXPHOS signature (Fig. 2C-D, Supplementary Table S7).

For characterization of the cellular components of the tumor micro-environment in each GBM subclass, we inferred the fraction of cellular states and stromal/immune cells with GBMdecon-voluteR^25^ and tumor cell purity by applying PUREE^37^. PN tumors were enriched in neural progenitor-like (NPC) cells, whereas PN-L tumors were associated with the presence of oligodendrocytes. CL-A tumors were enriched in astrocyte-like cells (AC). The CL-B subtype had a strong activated dendritic cell gene signature, suggesting that this subtype could potentially be treated with dendritic cell vaccines. CL-C tumors were enriched in mast cells. MES tumors displayed infiltration with macrophages, monocytes, and other immune cells. The NL subclass was associated with the presence of B cells, consistent with previous findings^22^, suggesting a distinctive immunological profile for NL gliomas (Fig. 2D). Tumor cell purity was lowest for NL, followed by MES and CL-B tumors, and tumor cell purity was highest for PN, CL-A and CL-C tumors (Fig. 2D).

We assessed the impact of our refined GBM classification on clinical outcomes for each of the subgroups, by using log-rank tests to compare the results obtained with those for Verhaak’s classification (Fig. 2F). Overall survival was highest for the PN-L subclass (Fig. 2F, *p*<0.001). By contrast, the worst outcomes were observed for the CL-C, MES and CL-A subclasses. Verhaak and Iavarone classified the CL-B and CL-C subclasses as CL and MTC, respectively, but our subdivision highlighted a clear difference in overall survival between these two subclasses. We also confirmed the reported associations with sex and age (Fig. 2G, Supplementary Table S8), including the overrepresentation of MES subclass tumors among samples from male patients and the lower median age of patients for the PN and PN-L subclasses than for the other subclasses. Interestingly, the CL-A group displayed no sex-related differences with respect to the other subgroups.

### Functional validation of Subtypes-Specific Coregulated Subnetworks through DepMap database

We investigated the subtype-specific active coregulatory networks by first inferring differentially Influencing regulators (DIRs) with the Seurat::findmarker function^38^ and the default Wilcoxon rank-sum test (Fig. 3A, Supplementary Table S9). We then used publicly available data^39^ regarding the effect of CRISPR-Cas9-mediated knockout (AVANA 19Q4 DepMap data) of the selected regulators in our defined GBM-derived cell lines for each subclass (Fig. 3B, Supplementary Table S9). As a starting step, we showed that we could employ CoRegQuery to assign a specific class with high confidence (class probability > 0.75) and robustness (using various transcriptomic profiling techniques, including microarray and RNA-seq sourced from six different datasets) to 33 of the 42 GBM cancer cell lines (GBCCL), thus revealing a heterogeneous pattern of subclasses across the GBM cell-line panel (Supplementary Table S10). This analysis also showed that GBM cell lines can be subtyped in 5 out of 7 classes, with the exception of the PN-L and the NL ones. Thus, we could not integrate the CRISPR-Cas9-mediated knockout data for these subtypes. In clinical samples, the PN coregulatory subnetwork confirmed the highly proliferative state of these tumors, visible on enrichment in several regulators predominantly involved in cell division, cell cycle, and proliferation such *CDC6, FOXM1, FOSL1, BIRC5, E2F7, PA2G4, DEPDC1, ATAD2*, and *TFAP2C.* In parallel, CRISPR-Cas9 AVANA 19Q4 analysis of the PN cell lines revealed a strong dependence on *BRCA2* and *FOXM1*, and a moderate but significant dependence on *GLI2*, and *PBX4* (Fig. 3C and Suppl. Table S10). The highest dependence of this subtype was for *PRMT5,* which appears as a strong regulator in this subtype and in PN-L and NL subtypes too. The PN-L class of tumors was characterized by high levels of activity of TFs showing high influence in the PN and the NL subtypes too, such as *SALL2, ZNF704, CAMK4*, and *CREB3L4,* but also by the uniquely high activity of *GBX2*, the Gastrulation Brain homeobox 2, recently described to act as a tumor suppressor in GBM cells, and inversely correlated to GBM grade^40^. This is in agreement with the most favorable clinical outcome shown by PN-L subtype of GBM. The NL coregulated subnetwork comprised the *PRDM16, SP100, TNIP1, ZFP3, MEIS3, POU3F3, SNCA, SMAD3 and TRB23* regulators. The products of genes such as *PRDM16* and *MEIS3* are instrumental in cell-lineage development and fate decisions. *PRDM16*, in particular, is linked to stem-cell maintenance and cell differentiation^41^. *SP100* and *SNCA* are correlated with lower levels of malignancy in GBM cell lines^42,43^ suggesting a potential role in mitigating malignancy in tumors of this class. Moreover, in the NL class, *SMAD3* has emerged as a regulator associated with favorable prognosis. Its inhibition, coupled with stimulation of the neurogliomal circuit, accelerates tumor progression^44^. Unsurprisingly, *SNAI2, PAX8, FOSL2, EPAS1, RUNX2,* and *EGFR* were associated not only with the top-ranking influence scores for MES-specific regulators, but also with the top dependency scores in MES GBM cell lines, highlighting their known roles in the mesenchymal state. In addition, our coregulated subnetwork for MES tumors confirmed the established role of the hypoxic tumor microenvironment, as suggested by the inclusion of *EPAS1* and *CEBPD*, both master transcription factors for hypoxia-regulated genes in GBM^45,46^. CRISPR-Cas9 AVANA 19Q4 analysis of the MES cell lines also revealed novel genes with a high influence and a strong dependence score, such as *ARHGEF5*, encoding for the Rho guanine nucleotide exchange factor 5, involved in the EMT process in other types of tumors^47^, but never studied in GBM.

**Figure 3.**
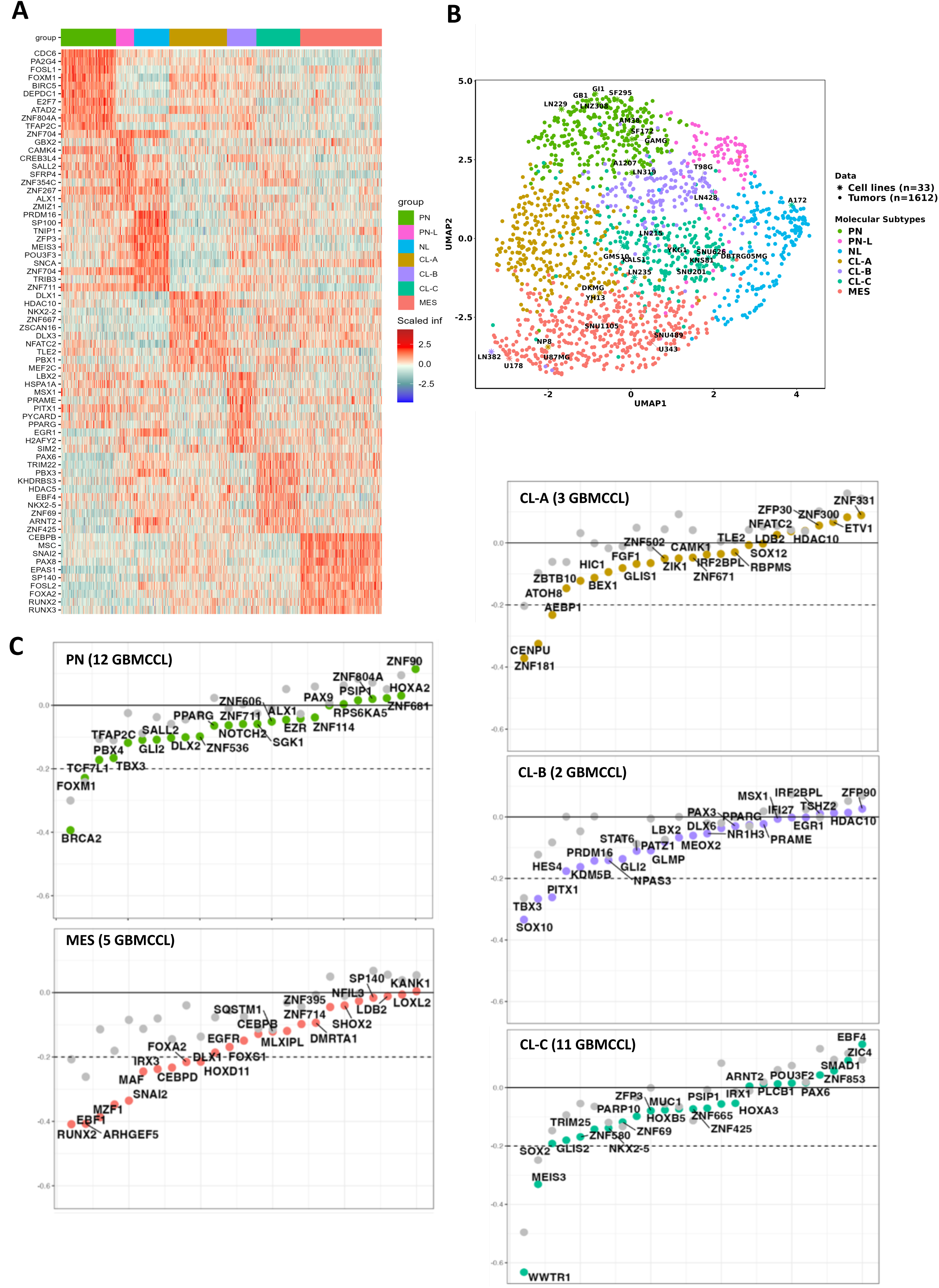
Identification of GBM subclass-matched cell-line repertoires and functional validation of specific differential subnetworks. (A) Heatmap showing the top 10 differentially Influencing regulators for each subclass. (B) Colocalization of the 42 GBM cancer cell lines on GBM-CoRegMap, as predicted by CoRegQuery, with labeling (color) according to the SVM-based classifier. (C) Impact on cell viability (CERES dependency score) for each of the predicted subclasses. For each regulator, two values are shown vertically: the colored dot is the mean CERES score for the cell lines assigned to the subclass, and the gray dot is the mean CERES score for cell lines not assigned to the subclass. A gene may have a negative CERES score (indicating that its knockout would reduce the survival of the corresponding cell line) or a positive score (indicating that its deletion might increase the survival of the cell line). The dotted horizontal line indicates the viability threshold used.

By contrast to the PN/MES axis, few TFs/co-TFs have been characterized so far as key regulators of Verhaak Classical tumors. Our analysis revealed that the CL-A coregulated subnetwork included epigenetic modifiers, such as *DLX1* and *HDAC10*, as well as *NKX2-2* and *TLE2* (Fig. 3A, Supplementary Table S10). However, the CRISPR-Cas9 analysis of CL-A cell lines uncovered a significant dependence on *ZNF181*, also showing a high influence score in CL-A samples, and of *CENPU*, encoding for the centromere-associated protein U, whose role has never been described in GBM so far (Fig. 3C). The CL-B coregulated subnetwork included *PRAME, PPARG, SALL1*, and *STAT6.* CRISPR-Cas9 analysis confirmed that the main regulators of CL-B cell lines are *PITX1*, *HES4,* and *SOX10*. Notably, the CL-B tumor subclass displays high levels of activity for the transcription factor *PPARG*, a master metabolic regulator implicated in tumor stromal-epithelial crosstalk and carcinogenesis^48^. Furthermore, the CL-B class relies heavily on the *PRAME* regulator (Fig. 3C), an oncoprotein linked to advanced tumor stages and poor prognosis, highlighting its clinical relevance^49^. Both *PPARG* and *PRAME* are among the TFs listed in the results of CRISPR-Cas9 screening in CL-B cell lines, further confirming the relevance of the data we obtained in the in vitro models (GBCCL), and its overlap with clinical sample results. The CL-C coregulated subnetwork included *PAX6, HDAC5, POU3F2, SOX2, NKX2-5, EBF4, MEIS3*, and *GLIS2.* In CL-C cell lines, the CRISPR-Cas9 analysis (Fig. 3C) indicated *SOX2*, *GLIS2*, and *NKX2.5* as deeply involved in survival, together with *WWTR1* and *MEIS3.* While the involvement of the core stem cell transcription factor *SOX2*^50,51^ and of the well known regulator of stemness and cell plasticity *WWTR1* are expected findings, our analysis uncovered the unprecedented importance of *NKX2.5*, *MEIS3*, and *GLIS2* in this subclass.

Note that the GBM-cRegMap tool allows the user to visualize active coregulated subnetworks and to explore any gene viability score, while also providing the calculated influence scores. As a representative example of this application, Fig. 4 depicts the MES coregulated subnetwork, where the *SNAI2* TF stands out for its influence, and, on the bottom, the influence and dependency score maps of *SNAI2* in the GBM cell lines.

**Figure 4.**
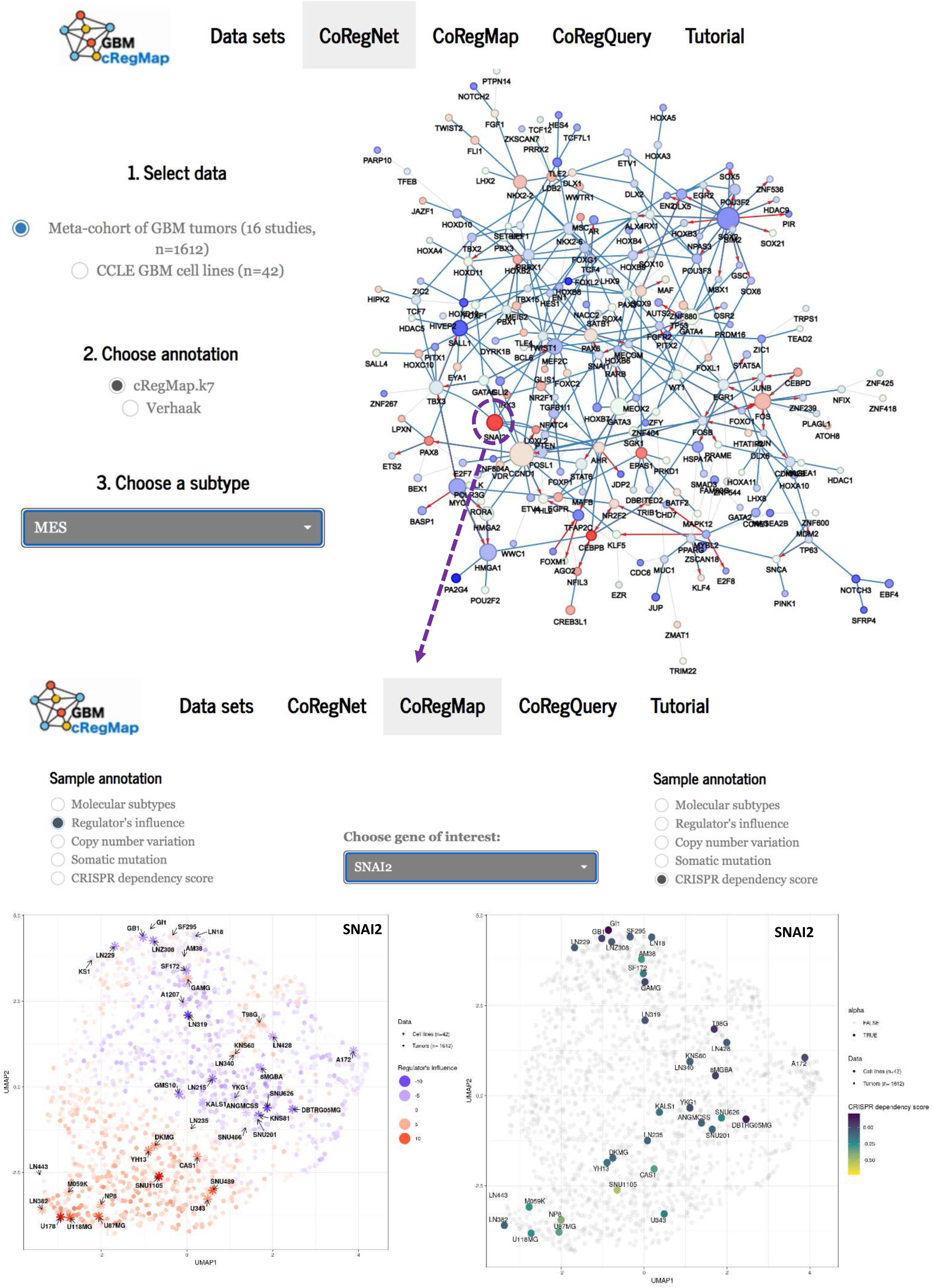
Example of exploration of MES co-regulated subnetwork using GBM-cRegMap synergistic compounds. The CoRegNet tab allows users to investigate the extent to which a phenotype of interest (e.g., MES subtypes), characterized by a set of active coregulators, colocalizes within the reference GBM-CoRegNet network through an intuitive visualization. Node color (red = high; blue = low) indicates the influence of the corresponding TF/coTF, with color intensity representing the strength of the influence. Node size and edge color follow the same patterns as indicated in Fig. 1A. The selected coregulators will then be analyzed sequentially, using the GBM-CoRegMap tab. The left plot displays the regulatory influence of the SNAI2 TF, while the right plot shows CRISPR dependency scores for SNAI2.

**Figure 5.**
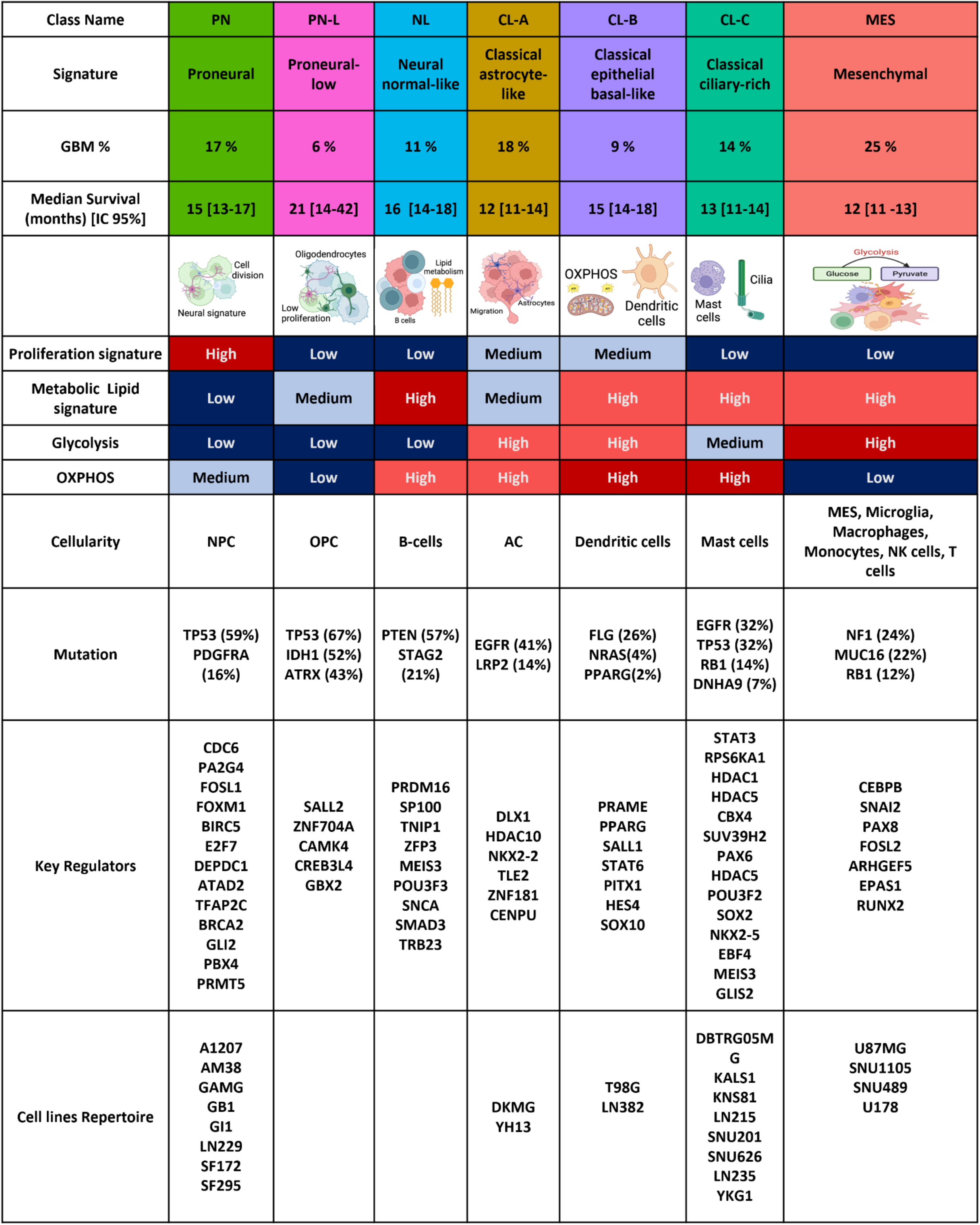
Summary of the main characteristics of the GBM-cRegMap classes. From top to bottom, the following information is displayed: Class abbreviation and name; proportion of classes in the 1612 samples of the metacohort; graphical representation of tumor cells and their characteristics; proliferation-based color scale, and a table showing, for each class, the most important characteristics such as active regulators, overexpressed gene markers, molecular alterations, enriched metabolic pathways, single cell state, immune cell infiltration, and median survival.

### Uncovering intermediate states in the PN-MES axis involved in chemotherapy resistance

Recent data have demonstrated the high degree of plasticity of GBM cells along a PN-MES axis^52^. Transitions between cellular states include intermediate states especially during therapies^53^. We applied GBM-CoRegQuery to analyze transcriptomic data (GSE253458: n = 9 samples), focusing on the comparison between the untreated U87MG glioma cell line and its TMZ-resistant variants, U87MG-R50 (continuously exposed to 50 µM TMZ) and U87MG-OFF (representing the post-TMZ removal condition)^54^. Unlike the Verhaak classification, the three cell lines were classified into three clearly distinct GBM-cRegMap subclasses (Fig. 6A, Supplementary Table S11): U87MG to the MES subclass (as already described), U87MG-R50 to the CL-C subclass, and U87MG-OFF to the PN subclass. This categorization highlights the dynamic transcriptional change of GBM cells’ behavior in response to chemotherapy and subsequent withdrawal of treatment, suggesting that CL-C subclass may represent an intermediate state in the PN-MES axis. The classification of U87MG-R50 within the CL-C subclass, characterized by a low proliferative signature, aligns with our prior in vitro studies, showing that U87MG-R50 exhibits a reduced proliferation rate compared to both U87MG and U87MG-OFF^54^. A crucial aspect of our analysis was the identification of regulators involved in the resistant states (Fig. 6B). In response to TMZ treatment and acquired resistance, the differentially activated subnetwork included MDM2 TF activation and corroborates our prior research on MDM2’s involvement in TMZ resistance via the p53 pathway^54^. Focusing on U87MG-R50/CL-C subclass hallmarks, the influence of *PAX8* was reduced while that of *NKX2.5* grew in these cells, compared with non-treated U87MG. In the U87MG-OFF cells, characterized by TMZ resistance after treatment removal, we observed a tendency of both *PAX8* and *NKX2.5* influence to return to untreated U87MG levels (Fig. 6C). Interestingly, the mRNA expression level of these regulators from the transcriptomic data did not significantly differ among the three cell lines. Thus, measuring of the influence effect of master regulators provides a new level of data interpretation. In line with this, we could experimentally confirm that PAX8 protein expression levels aligned with its predicted influence (Fig. 7A). Although very few explored, *PAX8* is detected in high-grade glioma and participates to glioma cell survival^55^ and is related with epithelial to mesenchymal transition (EMT) in other cancers^56^. *NKX2.5* is a transcription factor essentially involved in cardiac development^57^, and its expression/function has never been described in glioblastoma so far. Interestingly, when we analyzed its mRNA expression levels in the CGGA (Chinese Glioma Genome Atlas) cohort^58^ of primary (n = 651) and recurrent (n = 332) gliomas, we found that *NKX2.5* is significantly more expressed in grade IV gliomas compared to both grade III and grade II, and this difference holds true not only in primary but also in recurrent tumors (Fig. 7B). In the latter ones, a higher expression of *NKX2.5* negatively correlates with patients’ survival, even when restricted to recurrent GBM. (Fig 7B). This is particularly intriguing in view of the formation of temozolomide resistant, aggressive tumors after long-term treatments with this drug. Gene ontology (GO) enrichment analysis of the 282 *NKX2.5* putative target genes (175 activated and 107 repressed) revealed that *NKX2.5* negatively regulates genes implicated in glucose metabolism/glucose catabolic process to pyruvate (*e.g.,* HK2, GPI, GAPDH, LDHA), (Fig. 7C, Supplementary Table S12). Consequently, we experimentally verified the impact of *NKX2.5* overexpression in U87MG cells. A statistically significant decrease of all 4 genes mRNA was obtained accompanied by a decrease of the corresponding HK2 protein for example (Fig. 7C). *NKX2.5* was thus revealed through CoRegMap to be an interesting player in metabolic reprogramming in glioblastoma which deserves further studies.

**Figure 6.**
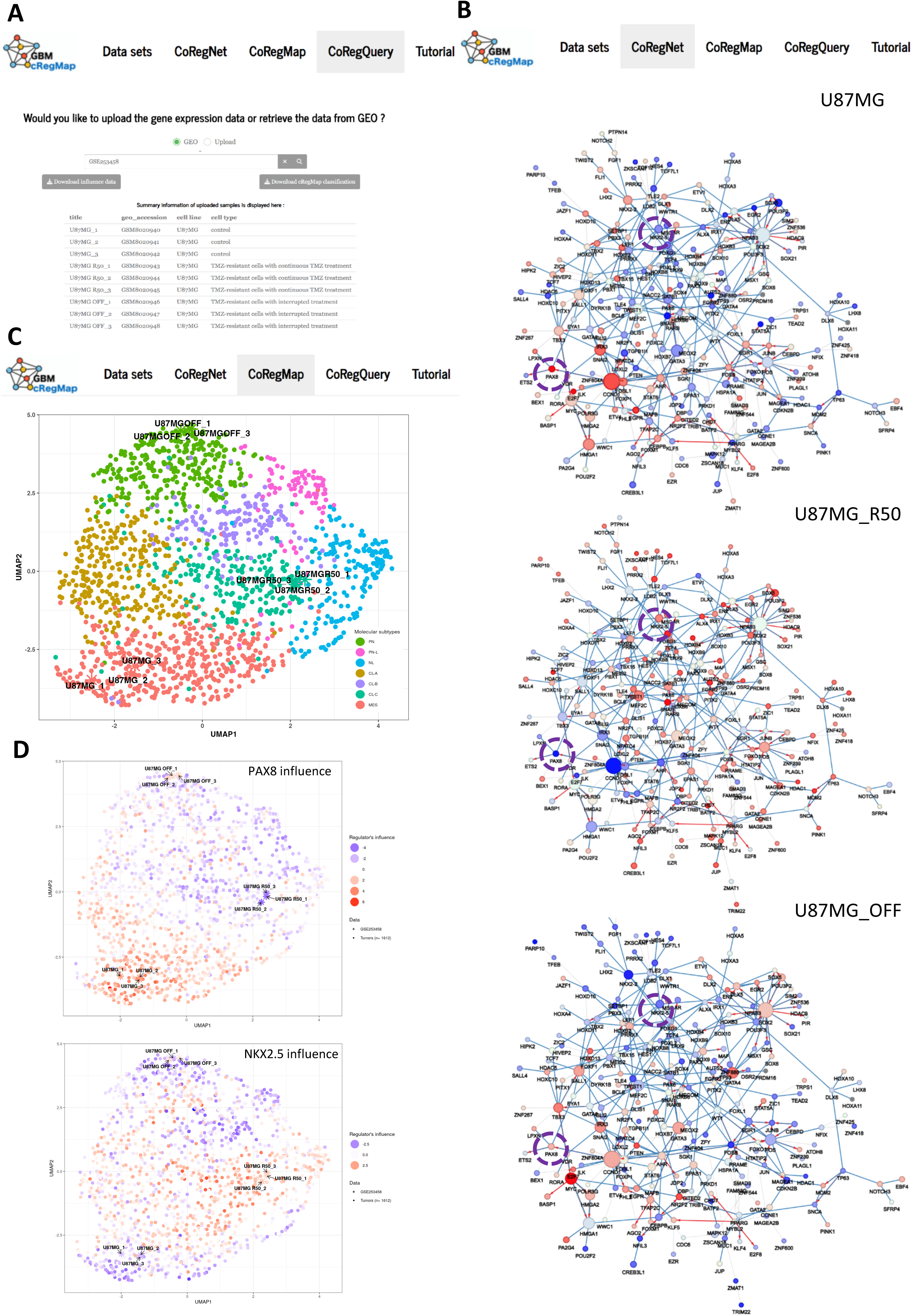
Example of reference-mapping and annotation of query dataset using CoRegQuery. (A) As input, users upload or simply provide a GEO series ID (GSE253458) of the query data using CoRegQuery tab (B) The CoRegNet tab allows users to investigate the extent to which studied cells (three replicates each) for naive U87MG cells and TMZ-resistant cells, U87MG-R50 and U87MG-OFF, are characterized by a set of active coregulators. Node size and color follow the same patterns as indicated in Fig. 4. (C) The GBM-CoRegMap tab provides colocalization information for the nine cell lines, as predicted by CoRegQuery, with labeling (color) according to the SVM-based classifier. (D) The top plot displays the regulatory influence of the PAX8 TF, while the bottom panel shows the regulatory influence of the NKX2.5 TF.

**Figure 7.**
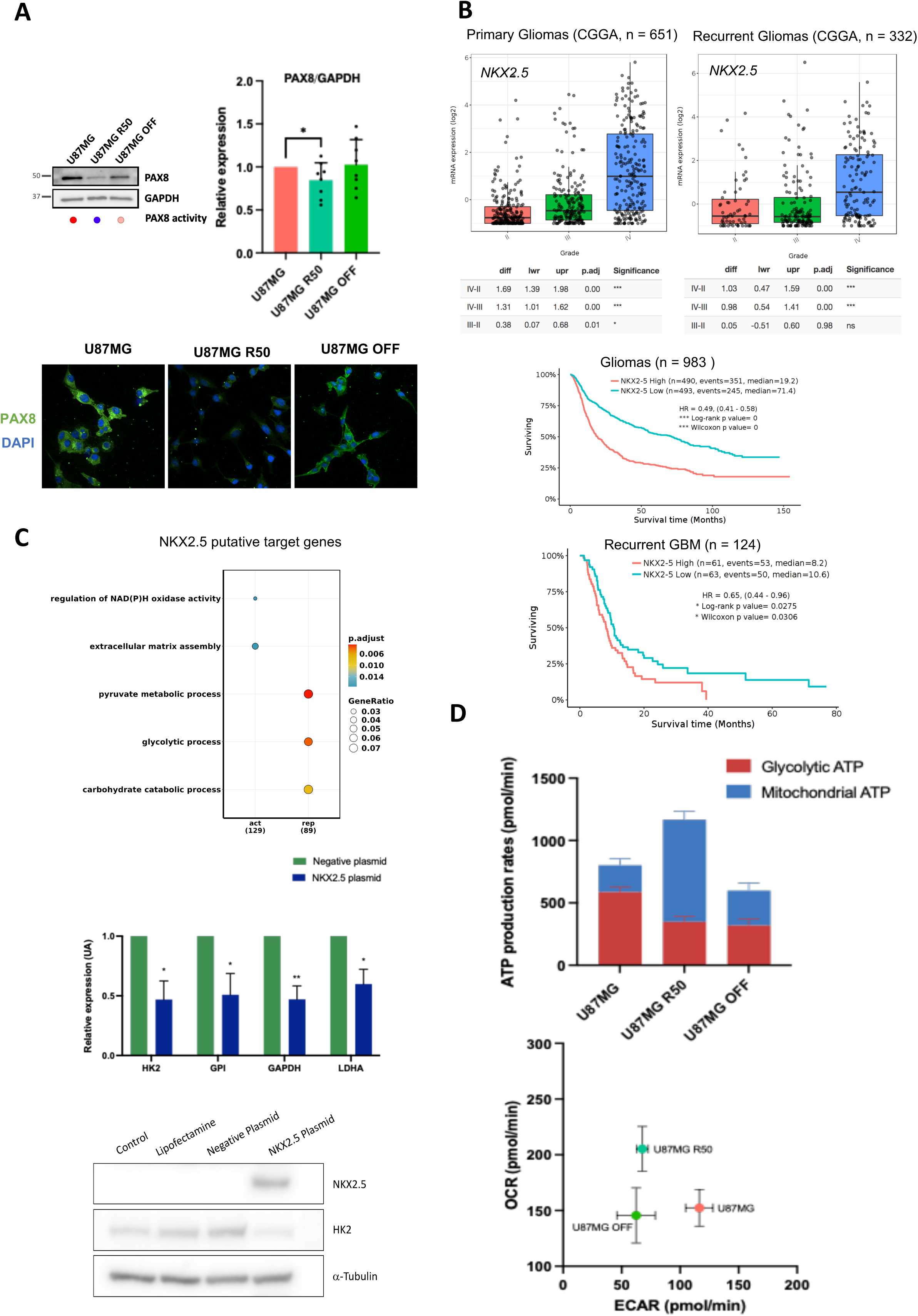
Functional validation of predicted phenotypic plasticity upon chemotherapy. (A) Analysis of PAX8 Protein Expression in U87MG, U87MG-R50, and U87MG-OFF Cells Representative Western blot detection of PAX8 protein and its predicted activity based on the regulatory network (colored dots under the WB), along with quantitative analysis (n>3) of its relative protein level (Top panel). Bottom panel: detection of the PAX8 protein by immunofluorescence in the same samples analyzed by WB. (B) Top panel shows the mRNA expression levels of NKX2.5 in primary and recurrent gliomas, as analyzed using the CGGA (Chinese Glioma Genome Atlas, http://www.cgga.org.cn/index.jsp) cohort. NKX2.5 expression is significantly higher in grade IV gliomas compared to grade III and grade II gliomas. Bottom panel illustrates the survival analysis of recurrent GBM patients (grade IV) based on NKX2.5 expression levels. (C) *NKX2.5* Mediated Transcriptional Repression of Glycolytic Genes. Top Panel: Functional enrichment analysis of the repressed and activated genes of NKX2.5 using clusterProfiler. The plot visualizes the enriched Gene Ontology Biological Processes (GO-BP) terms, with the size of each dot representing the gene count and the color indicating the adjusted p-value, reflecting the significance of the enrichment. Middle Panel: RT-qPCR analysis showing the reduced expression of glycolytic genes (HK2, GPI, GAPDH, LDHA) in U87MG cells upon overexpression of NKX2.5. Bottom Panel: Western Blot Analysis of NKX2.5 and HK2 protein expression in U87MG cells in 4 conditions: control (untreated cells), lipofectamine treated cells, negative control plasmid and NKX2.5 plasmid transfected cells. (D) Metabolic signatures of assigned classes for each cell line (U87MG, U87MG-R50, and U87MG-OFF) and energy metabolism characterization using Seahorse XFp extracellular flux analysis. This includes ATP production rates and the contribution of glycolysis and OXPHOS, as well as an energy map, depicting both Oxygen Consumption Rates (OCR) and Extracellular Acidification Rates (ECAR). All Seahorse data are represented as means from three independent experiments ± SEM.

Furthermore, the metabolic profiles of our cell lines, as determined by extracellular flux analysis, were consistent with the metabolic signatures of their assigned subclasses (Fig. 7D). U87MG displayed a predominantly glycolytic profile, aligning with the glycolytic signature of the MES class. U87MG-R50 exhibited an OXPHOS-dominant profile, in line with the metabolic characteristics of the CL-C subclass. In contrast, U87MG-OFF was metabolically less active, reflecting the repressed metabolic signature associated with the PN subclass. Taking together, these results confirm the CL-C class represented by U87MG R50 as an important intermediate class in the response to therapies and shows that the associated transcriptomic program can be easily modeled in co-regulatory network, which is based on TF/coTFs influence and can therefore be used to identify putative master regulators of biological traits.

## Discussion

We have developed GBM-cRegMap, a powerful web-based tool, to provide researchers with rapid access to a unified coregulatory influence network view of GBM. Many of the software tools used are based on techniques originating from artificial intelligence (AI), especially representation learning: h-LICORN^9^, CoRegNet^13^, LatNet^59^ and Seurat^38^ (Graphical abstract). We describe here the first ever use of transcriptomes from GBM tumor-derived cell lines to produce a more reliable GBM-CoRegNet and to prevent the introduction of possible bias due to the use of less homogeneous transcriptomic data from GBM tumors presenting non-specific signals and signals associated with the tumor microenvironment. We then demonstrated the relevance of building a harmonized tumor heterogeneity map, GBM-CoRegMap, encompassing a meta-cohort analysis of 1612 tumors from 16 studies. We demonstrated that GBM-CoRegMap successfully removed batch effects and reproduced the known heterogeneity of GBM. More importantly, our comprehensive GBM-CoRegMap gave rise to a novel molecular classification of GBM into seven distinct classes. Each subtype is uniquely defined by the activity of a regulatory network with distinct biological and clinical implications (Fig. 5). We confirmed and expanded our knowledge of the proneural (PN) and mesenchymal (MES) subtypes and distinguished three classical-like subtypes (CL-A: astrocyte-like, CL-B: epithelial basal-like, CL-C: cilium-rich), in addition to NL (neural normal-like) and weakly proliferating PN-L (proneural-Low) subtypes. The discovery of different subtypes of classical GBM, such as CL-C, rich in ciliary features, and CL-B, with epithelial traits, has highlighted the roles of unique molecular and biological processes. The poor prognosis of the CL-C subtype, linked to its ciliary characteristics, highlights the potential role of primary cilia in GBM oncogenesis and resistance to treatment. This finding suggests that the targeting of cilium-associated pathways may be crucial, particularly for the treatment of challenging subtypes, such as CL-C. In this regard, the influence and dependence score of GLIS2 in the CL-C subtype is particularly interesting, as this protein is a transcriptional repressor which localizes at the ciliary base, and works a central regulator of stem and progenitor cell maintenance and differentiation^60^. Our findings provide evidence in support of the existence of a normal-like (NL) subtype, as identified by Nguyen *et al.,* in 2020^22^, corresponding to 49% of the NEU_Verhaak subtype tumors. The NL subtype is characterized by a “normal” transcriptomic profile enriched in neuronal development-associated pathways. There is increasing evidence to suggest that some GBMs hijack neuronal mechanisms to fuel their own growth^55^. This finding challenges the view put forward by Wang et al.^4^, who suggested that the neural subtype defined by Verhaak might lack a tumor-specific signature, thereby calling into question the identification of neural tumors as a distinct tumor subtype. We also discovered alterations to the tumor suppressor gene *PTEN* in 57% of NL cases, together with a higher frequency of *STAG2* mutations in this subtype, highlighting the distinctive features of these tumors, and possibly indicating a therapeutic window specific for this subtype.

An additional achievement of our work, based on our analysis of the influence and dependence scores of TFs and coTFs, is the identification of key regulators for each subclass. These factors not only represent those which most strongly and widely regulate the gene expression typical of each subtype, but also those whose depletion mostly affects the survival of GBM cell lines belonging to a specific subtype. This, in turn, has the intriguing implication that they might be explored as potential subtype-specific targets. The key regulators of the PN subclass were *ZNF536, BRCA2, GLI2, PBX4*, and *PRMT5.* Notably, *ZNF536,* previously described as a repressor of neuron differentiation^61^, has never been involved in glioma development. The very strong dependence score of the PN subtype on PRMT5 is in agreement with and integrates the findings of Sachamitr *et al*.^62^ describing that PRMT5 is a key target for the inhibition of GBM proneural subtype. The high influence and dependence score assigned to PITX1 in the CL-B subtype may point to a particular tendency of this subtype to set up neurogliomal synaptic communications, where the contribution of *PITX1* was recently demonstrated^44^. Even in the already well characterized MES subtype, we were able to confirm known regulators and a specific vulnerability, such as that on *RUNX2* expression^63^, and we also uncovered a novel one, depending on *ARHGEF5*.

More interestingly, the CoRegQuery tab maps new data to a reliable tissue-specific co-regulatory network and a large-scale manifold of tumor heterogeneity references. Reference-based analysis shifts data interpretation from unsupervised to supervised, allowing accumulated information from prior experiments to aid in interpreting new data. The here-described use case of GBM cell phenotypic plasticity in response to TMZ treatment demonstrates the potential of the CoRegQuery algorithm to enhance the analysis and interpretation of small transcriptomic datasets. Our analysis reveals the CL-C subclass, an intermediate state demonstrating the plasticity of GBM cells along the PN-MES axis under chemotherapy. We identified key regulators, such as MDM2, PAX8, and NKX2.5, involved in TMZ resistance. Notably, NKX2.5, more expressed in higher-grade gliomas, negatively impacts patient survival and repress genes involved in glucose metabolism. These findings suggest that the CL-C subclass is crucial in therapy response, providing insights into glioblastoma treatment resistance and identifying potential targets for future research.

In conclusion, we provide a comprehensive new tool for use in studies of the co-regulatory networks driving the heterogeneity and plasticity of GBM. This offers great promise for the GBM clinical and research communities to better understand the complex landscape of GBM and opens up new possibilities for characterizing key regulators and suitable therapeutic targets according to the molecular heterogeneity and plasticity of GBM.

## Methods

### Public transcriptome data collection

Transcriptomic and clinical outcome data for GBM tumors were collected from GlioVis^66^, including 16 datasets corresponding to 1612 samples (Supplementary Table S1). Additional transcriptomic data for cell lines were downloaded from the Cancer Dependency Map (DepMap)^39^ and GEO, including 10 datasets corresponding to 42 commercial GBM cell lines, and eight normal brain samples (Supplementary Tables S1, S11 and S12). For consistency, all gene identifiers were transformed into HUGO Gene Nomenclature Committee (HGNC) symbols. All the above datasets were also classified using the Verhaak classifier from the GlioVis web application. The RNA-Seq datasets that had been downloaded from the GEO database were normalized with the edgeR^67^ R package, with TMM normalization and log-transformed counts per million. Microarray data were normalized with the standard workflow of the limma^68^ R package.

### Public data collection for regulatory elements and interactions

A list of regulators corresponding to 2,375 genes from two different sources (1,639 transcription factors (TF) with experimentally validated DNA binding specificity^69^ and 752 transcription cofactors (TcoF) with experimental validation information^70^) was used. The regulation evidence (TF-Target) list included 1,107,092 interactions retrieved with a wide range of bioinformatics tools and data-bases (listed in Supplementary Table S2) from 1) ChIP-seq data downloaded directly from the ChEA2 database and 2) human transcription factor binding site (TFBS) models, in the form of position weight matrices (PWM) recovered with MotifDB R/Bioconductor. The promoter sequences were scanned for these TFBS with the PWMEnrich R/Bioconductor package, and 3) completion of the TF-Target list with the tftargets R package and evidence from databases (*e.g.,* TRED, ITFP, ENCODE, BEDOPS, TRRUST). The co-operative evidence (TF-TF) list contained 1,149,582 protein-protein interactions (PPI) from various PPI databases (*e.g.,* BioGrid, HIPPIE, STRING, and HPRD) obtained with the iRefR R package.

### Gene regulatory network inference

Gene regulatory networks were inferred with the CoRegNet Bioconductor package^13^. Starting from transcriptomic data and a list of human regulators, CoRegNet uses the hLICORN algorithm^9^ to capture regulatory interactions between regulators and target genes in four steps: (1) Transcriptomic data is discretized into values of −1, 0 and 1 according to per-gene distribution. Genes present in the transcriptomic dataset were split into regulators and target genes, and only those with minimum support for non-zero values after discretization were retained. (2) A frequent itemset algorithm identifies potential sets of co-activators and co-inhibitors from the list of regulators. (3) For each target gene, a list of candidate co-activator and co-inhibitor sets (GRN) was selected according to a regulatory program (association rule) metric. (4) Each GRN candidate for each gene was scored based on the regression between the expression of the GRN set regulators and the expression of the gene concerned. For each gene, we retained the top ten GRN candidates according to R2 score. CoRegNet can then be used to select the best GRN for each gene based on the interaction evidence data. Evidence of regulation and cooperation was incorporated into each candidate GRN with an integrative selection algorithm, yielding an R2 score for each of the integrated datasets. The GRN with the highest merged score was selected. CoRegNet builds a coregulatory network, GBM-CoRegNet, from the GRNs obtained, by setting a cooperativity relationship between a pair of regulators, TFi and TFj, if they have a minimum of five target genes in common and the relationship is significant (*p* =< 0.01) according to the Jaccard similarity co-efficient formula: 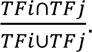

### Network-based regulatory influence signal quantification

The regulatory network structure provides a set of genes that are activated and repressed under reference conditions. Based on this structure, we can capture the influence of each regulator, a latent signal of the regulator activity in each sample based on its observed effect on downstream entities. For each regulator, Welch’s *t*-test was performed to compare the distribution of activated (Ar) and repressed (Ir) genes. The influence of a regulator *r* is calculated as follows:

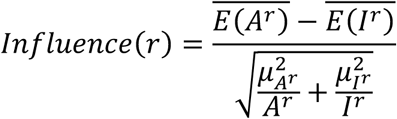

where *E(A^r^)* and *E(I^r^)* are the expression of the activated and repressed genes in the samples, respectively. 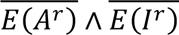 are their respective means, and *μA^r^* and *μI^r^* are their standard deviation. A regulator is active only when it activates A^r^ and represses I^r^ in accordance with the expectations of the regulatory network model, giving a positive *t* value in the Welch *t*-test. Higher *Influence*(*r*) values indicate higher levels of regulator *r* activity in the sample. In this work, we calculated the *Influence* of a regulator only if it has activated or repressed at least five genes. Due to the strict thresholds used in network construction, some local gene networks may be filtered out, resulting in some regulators having activated or repressed target gene sets with fewer than five members. In such cases, we fill the smaller set (fewer than 5 target genes) with the target genes for which the regulator concerned has the highest R2 regression score. This makes it possible to capture the *Influence* values for a maximum number of regulators. In addition, for some regulators, expression values may be missing for some of the target genes in some tumor samples for which we wish to assess the influence. In this case, the missing expression values are estimated with the LatNet method^59^, using expression levels in cell lines as a reference dataset. It may not be possible to estimate the expression of some target genes for a particular sample, and this may preclude calculation of the influence of a particular regulator. In this (rare) case, the median of the obtained influence values for the regulator in the other samples is used to replace the missing value.

### Seurat clustering

cRegMap subclass computation was performed with the Seurat R package^38^, by a modularity optimization-based clustering approach. The smart local moving (SLM) graph clustering algorithm was applied to a shared nearest neighbor graph (SNN) plotted from the influence data. Before clustering with the Seurat package, the Seurat:: ScaleData function was used with default parameters to scale the influence data. Seurat::RunPCA results were called with 20 PCs, and all other parameters were set at default values. Clustering with the Seurat package was performed on all calculated principal components with the Seurat::Find-Neighbors function with default parameters, followed by the Seurat::FindClusters function with a specified resolution of 0.8, resulting in seven clusters.

### Reference-mapping and annotating query datasets

Query datasets are mapped and annotated in a multistep process. CoRegQuery first uses GBM-CoRegNet to compute the regulatory influences of the input dataset. A support vector machine (SVM) model is then used to classify the query samples. We used the e1071 R package^71^, implementing an SVM with a radial kernel and a one-vs.-rest strategy for multiclass classification. The SVM model was trained on the influences and classifications derived from the tumor cohort. In the next step, CoRegQuery uses the Seurat::ProjectUMAP function to project the query dataset samples onto the UMAP (uniform manifold approximation and projection) coordinates of GBM-CoReg-Map. The Seurat function identifies the nearest reference sample neighbors and their distances for each query sample and performs a UMAP projection by supplying the calculated neighbor set along with the UMAP model previously computed in GBM-CoRegMap.

### Functional enrichment analysis of DEGs

For microarray gene expression analysis, the linear models for microarray data (LIMMA) R package^68^ was used, and *P*-values were adjusted by the Benjamini-Hochberg method. For RNA-Seq data gene expression analysis, the edgeR^67^ R package was used, followed by limma-voom^72^ and standard differential expression (DE) analysis. Gene Ontology (GO) enrichment analysis was performed with the R packages clusterProfiler^73^ and enrichR^74^. Ontologies with a *p_adj_*<0.01 and gene counts of more than five were examined. The Kyoto Encyclopedia of Genes and Genomes (KEGG), Molecular Signatures Database (MSigDB), and REACTOME pathway databases were searched for enriched terms with the R package msigdbr.

### Analysis of genomic alterations

Somatic mutations were analyzed with the GBM-TCGA dataset. The TCGAbiolinks R package was used to download mutation annotation files (MAF) aligned with the hg38 sequence, and analysis was performed with the R package maftools.

### Deconvolution of cellular components and tumor cell purity scores

The TCGA-GBM dataset was used to determine the tumor cell purity of each sample and the abundance of cell populations with respect to the GBM-cRegMap classification. The PUREE^37^ (pan-cancer tumor purity estimation) web server was used to calculate the purity score for each sample of the dataset and to display it as a boxplot for each class. The GBMdeconvoluteR^25^ web server was then used to estimate the abundance of various immune and stromal cell populations in each sample, and the resulting values were averaged by GBM-cRegMap subclass.

### Survival analysis

Survival analyses were performed with the R package survminer; overall survival was calculated by the Kaplan-Meier method, and log-rank tests were performed to compare survival curves.

### Classification of cell lines

CoRegQuery assigned a specific class with a high degree of confidence (SVM posterior probability > 0.75) to 33 of 42 GBM cancer cell lines (GBCCL) (Fig.4, Supplementary Table S11), revealing a heterogeneous pattern of subclasses across the GBM cell-line panel. We assessed the robustness of our method by analyzing multiple independent datasets for a single cell line. These datasets were generated with various transcriptomic profiling techniques, including microarrays and RNA-Seq, and were sourced from six different datasets (Supplementary Table S9). The classification outcomes were consistent across the different profiling techniques and datasets, demonstrating the reliability of our classification approach. An exploration of the PubMed citation status of the studied GBCCL (Supplementary Table S11) revealed a bias, with only five (150> citations each) cell lines used by the scientific community. Our classification system was consistent with published results for these five cell lines. For instance, U87MG, a well-known mesenchymal GBCCL, was assigned to the MES subclass, LN229 to the PN subclass, and T98G, which is known to be dependent on oxidative phosphorylation^75^, to the CL-B class. Normal brain samples (GSE15824, *n*=2; GSE15209, *n*=6) were categorized as NL.

### Identification and validation of subclass-specific regulators

We used Wilcoxon rank-sum tests to identify differentially Influencing regulators (DIRs) with the Seurat::FindAllMarkers function (min.pct= 0.25 and logFC.threshold= 0.20) and *p*-value adjustment by Bonferroni correction. Wilcoxon-Mann-Whitney tests between the mean CERES dependency score of the cell lines assigned to the subclass and the mean dependency score of the rest to rank the DIRs further. The CERES dependency score of the cell lines was acquired with the R package depmap^39^.

### Cell lines, culture, and conditions

The U87MG GBM cell line was obtained from the American Type Culture Collection (LGC Standards Sarl, Molsheim, France). Two TMZ-resistant clonal lines were established in our laboratory by exposing the parental U87MG line to 50 μM TMZ for extended periods: U87MG-R50 and U87MG-OFF. U87MG-R50 cells were continuously cultured in a medium supplemented with 50 μM TMZ, whereas U87MG R50 OFF cells were treated with TMZ for two months, after which they were grown in TMZ-free medium. TMZ (cat #T2577; Sigma Aldrich) was prepared as a 100 mM stock solution in DMSO and stored at 4°C until use. These cell lines were routinely cultured in Eagle’s minimum essential medium (EMEM) supplemented with 10% heat-inactivated fetal bovine serum (FBS), 1% sodium pyruvate, and 1% non-essential amino acids, at 37°C, under an atmosphere containing 5% CO2.

### Immunofluorescence assays

Cells were fixed by incubation with ice-cold methanol for 10 min. They were then incubated overnight at 4°C with the PAX8 Proteintech #10336-1-AP primary antibody (1:50). The cells were then incubated with a goat anti-rabbit AF488 secondary antibody (1:500) at room temperature for 2 h. The expression of PAX8 was assessed in U87MG, U87MG R50, and U87MG R50 OFF cell lines and analyzed under a fluorescence microscope.

### Seahorse metabolic flux analysis

Real-time analysis of oxygen consumption rates (OCR) and extracellular acidification rates (ECAR) was conducted using the Seahorse XFp analyzer (Agilent). Prior to analysis, XFp flux cartridges were pre-hydrated with XF Calibrant overnight at 37 °C. Microplates were coated with Cell-Tak at a concentration of 22.4 µg/mL (Corning®) to ensure cell adherence. Cells were seeded at a density of 60,000 cells per well in lineage-specific medium and both seeding, and analysis were performed on the same day. ATP production rates were determined using the Seahorse XF Real-Time ATP Rate Assay Kit, following the manufacturer’s recommended protocol using the medium recommended by the manufacturer (DMEM, 10 mM glucose, 2 mM glutamine, and 1 mM pyruvate, pH 7.4). OCR and ECAR were assessed under basal conditions and in response to sequential injections of Oligomycin (1.5 μM) and Rotenone/Antimycin A (0.5 μM each).

### Western blotting analysis

Cells were lysed in Laemmli sample buffer (Biorad) supplemented with 5% β-mercaptoethanol and proteins were fractionated by SDS-PAGE (20%), transferred to PVDF membrane (GE Healthcare, Velizy, France), which were blocked with 5% (w/v) nonfat milk in PBST for 1 h at room temperature, probed with primary antibody in blocking solution overnight at 4°C, incubated with horseradish peroxidase (HRP)-conjugated secondary antibodies (GE Healthcare) for 1 h at room temperature and the chemiluminescent signal was detected using enhanced chemiluminescence (ECL) system (ECL™ Prime Western Blotting System, GE Healthcare Bioscience) with a ImageQuant™ LAS 4000 analyser (GE Healthcare). Quantification of non-saturated images was done using ImageJ software (National institutes of Health, Bethesda, MD, USA). For each experiment, 3 lysates from different cell cultures were used. Tubulin or GAPDH were used as the loading control for all samples. The primary antibodies used were antibodies for NKX2-5 (Cell Signaling, cat #8792, 1:500), HK2 (Proteintech, cat #22029-1-AP, 1:5000) and α-Tubulin (cat #T9026, Sigma-Aldrich, 1:1000 dilution). The secondary antibodies used were anti-mouse IgG-HRP (cat #W4028, Promega, 1:10000 dilution), anti-rabbit IgG-HRP (cat#W4018, Promega, 1:10000 dilution).

### RT-qPCR analysis

U87MG/U87MG R50/U87 MG OFF were seeded in a 6-well plate at a density of 0.5 million cells per well and cultured in pure OptiMEM medium, or supplemented with Lipofectamine 2000 (ThermoFisher), negative control plasmid (5 µg), or NKX2.5 plasmid (5 µg) for a period of 24 hours. The cells were then washed with 1X PBS and lysed with 350 µL of RLT buffer from the extraction kit (RNEasy Plus Mini Kit, Qiagen) according to the manufacturer’s instructions. RNA concentrations were measured using an ND-1000 NanoDrop spectrophotometer (Labtech, Palaiseau, France). 1 µg of extracted RNA was used for cDNA synthesis using iScript Reverse Transcription SuperMix kit (BioRad). cDNA was analyzed by Fast SYBR™ Green (Applied Biosystem, Thermo Fisher Scientific, Massachusetts, USA) in duplicate, using the StepOne real-time PCR system (Applied Biosystem). qRT-PCR data were analyzed using StepOne Plus software. The relative expression of each target was calculated using the relative quantification method (2^-ΔΔCT^) with RNA18S (5’-TGTGGTGTTGAGGAAA-GCAG-3’ and 3’-TCCAGACCATTGGCTAGGAC-5’; Invitrogen) as internal control. GAPDH primers were 5’-GTCACCAGGGCTGCTTTTAACTCT-3’ and 3’-GGAATCATATTGGAACATGTAAACCAT-5’; Invitrogen). The following primer pairs were from Qiagen (RT2 qPCR primer assay): GPI (PPH00897C), HK2 (PPH00983B), LDHA (PPH02047H).

## Data availability

Public data used in this article were acquired from GEO (https://www.ncbi.nlm.nih.gov/geo/), DepMap (https://depmap.org/portal/download/all/), GlioVis (http://gliovis.bioinfo.cnio.es), and TCGA (https://portal.gdc.cancer.gov/). The in-house dataset of the U87MG cell line and its TMZ-resistant variants, which we made available, can be found in the GEO repository under accession number GSE253458.

## Code availability

GBM-cRegMap is written in the R language and uses the functions of several widely used packages (Supplementary Table S2) available from Bioconductor and CRAN, based on the Golem ^64^ framework. This framework simplifies the development and deployment of a stable and robust web application with R Shiny^65^. A plotly graphics system was used to generate interactive visualizations, with interactions enabled by brushing or clicking on them in the Shiny framework. The GBM-cRegMap has been released on Docker with all the necessary packages to prevent update conflicts. The GBM-cRegMap web application was deployed with Google Cloud Run and is publicly available from https://gbm.cregmap.com. The source code is available from GitHub (http://github.com/i3bionet/gbm-cregmap) under the Apache 2 Licence.

## Acknowledgments

This work was supported by grants from the *Institut National du Cancer* INCa (PL-Bio project no. 2017-145) and the French National Research Agency (ANR-20-CE20-0028). CB received a doctoral fellowship from the French Ministry of Higher Education, Research, and Innovation (MESRI). GP received a doctoral fellowship the French National Research Agency (ANR) AI_PhD@Lille program. MM received a doctoral fellowship from CNRS (MITI 80 PRIME program).

## Supplementary Tables Legends

**Table S1:** Transcriptomic datasets used to generate GBM-cRegMap reference compounds. Related to Graphical Abstract.

**Table S2:** Software, tools and resources required to develop GBM-cRegMap. Related to Graphical Abstract.

**Table S3:** GBM-CoRegNet regulators. Related to Fig. 1A.

**Table S4:** GBM-CoRegNet coregulator pairs. Related to Fig. 1A.

**Table S5:** Cross-study prediction performance. Related to Fig. 1D.

**Table S6:** Differentially expressed genes (DEGs) analysis. Related to Fig. 2C.

**Table S7:** Functional enrichment analysis for the GBM-cRegMap subclasses. Related to Fig.2C.

**Table S8:** Sex- and age-related differences in GBM-cRegMap subclasses. Related to Fig. 2G.

**Table S9:** Differentially Influencing regulators (DIRs) analysis. Related to Fig. 3A.

**Table S10:** Classification of GBM cell lines. Related to Fig. 3B.

**Table S11:** Classification of U87MG, and TMZ-resistant U87MG variants. Related to Fig. 6.C **Table S12**: Functional enrichment analysis (GO-BP) of the repressed and activated genes of NKX2.5 TF using clusterProfiler. Related to Fig. 7C.

